# YieldLearn: An interpretable machine-learning pipeline for estimating yield-related traits based on physiological indicators

**DOI:** 10.64898/2026.09.22.753232

**Authors:** Xi Zhang, Hamid Shaterian, Paula Ashe

## Abstract

Efficient yield prediction under field conditions remains a key challenge for crop breeders and agronomists. YieldLearn is an interpretable machine-learning pipeline developed to estimate yield-related traits in durum wheat (*Triticum durum*) using physiological indicators measured by the Hansatech Instruments Pocket PEA. The workflow integrates fluorescence parameters (Area, Fo, Fm, Fv/Fm, PI Inst.) with field-measured agronomic traits (Booting, Height, Maturity, Lodging, Yield kg/plot, and Adjusted Yield kg/ha). Five supervised learning algorithms multinomial logistic regression, random forest, support vector machine (radial kernel), k-nearest neighbors, and decision tree were evaluated using five-fold cross-validation. Models achieved average classification accuracy around 0.57 with AUC values exceeding 0.58. SHAP (SHapley Additive exPlanations) visualizations revealed that high Fv/Fm and PI Inst. values were positively associated with yield potential, whereas elevated Fo indicated stress-induced yield reduction. An interactive Shiny App enables real-time prediction and feature-importance exploration. Given the small sample size (165 plots) and absence of environmental covariates such as temperature or soil moisture, YieldLearn should be regarded as a reproducible exploratory framework rather than a production-ready predictor. Nonetheless, it offers a valuable foundation for guiding experimental design, validating physiological indicators, and supporting the next generation of scalable, interpretable crop-yield models.

## 1 Introduction

Accurate estimation of crop yield under variable environmental conditions is essential for advancing breeding programs and improving agronomic decision-making [1]. Traditional yield trials depend on destructive sampling and post-harvest analysis, limiting throughput and delaying feedback for selection or management. In contrast, chlorophyll fluorescence provides a rapid, non-invasive measure of photosynthetic efficiency and stress response, offering a potential physiological proxy for yield potential [2]. Recent genomic studies in barley have demonstrated that photosynthesis-related parameters such as ΦPSII and NPQt can capture heritable genetic variation relevant to yield formation under field conditions [3]. Furthermore, portable fluorometers such as the Hansatech Pocket PEA allow field-scale collection of photosystem II (PSII) parameters, including Fo, Fm, Fv/Fm, PI Inst., and Area, which together characterize photochemical performance and plant vitality. Despite this capability, most fluorescence-based yield studies rely on simple correlations or linear regressions that cannot fully capture the nonlinear relationships between physiological signals and yield. Recently, a group of researchers applied random forest and artificial neural network to predict the crop field yields with the plant fluorescence from satellite images [4]. But the reproducible, interpretable pipelines designed for agricultural datasets remain limited.

Recent advances in machine learning (ML) have transformed predictive modeling across many research areas [5]. Inspired by such approaches, we adapt this interpretability-driven methodology to crop phenotyping. This study introduces YieldLearn, a compact yet transparent ML workflow for fluorescence-based yield classification. The system integrates multiple supervised algorithms, applies SHAP analysis for feature attribution, and deploys an interactive Shiny App for real-time prediction and visualization. The objectives were to: 1. Evaluate the performance of multiple ML classifiers for predicting high vs low yield categories using fluorescence features; 2. Quantify the physiological contribution of each feature through SHAP interpretation; and 3. Provide an open, reproducible tool to support agronomic decision-making and future digital-agriculture research. The remainder of this paper describes the dataset and modeling pipeline, presents comparative results, and discusses implications, limitations, and opportunities for extension to larger, multi-environment trials.

## 2 Materials and Methods

### 2.1 Study context and data

The dataset originated from a field-trial experiment on durum wheat (Triticum durum) conducted by the personnels from the National Research Council Canada (NRC), Saskat oon. The physiological measurements were collected independently in the morning of mid-July and mid-August after three continuous sunny days. Each observation corresponds to a distinct experimental plot identified by *PlotID* and *LineID*. Physiological measurements were collected using the Hansatech Pocket PEA, capturing Fo, Fm, Fv/Fm, PI Inst., and Area—parameters describing PSII efficiency and cumulative electron-transport activity. Agronomic traits were measured separately from each *PlotID* including Booting (days), Height (cm), Maturity (days), Lodging (0-9 scale), Yield (kg/plot), and Adjusted Yield (kg/ha).

### 2.2 Data preparation

Fluorescence and yield data were merged by *PlotID*. Outliers were removed using interquartile filtering, and numeric predictors were standardized to unit variance. Binary yield classes (High/Low) were generated via median splitting, which is a robust approach for small datasets with micro-environmental variability.

### 2.3 Model training

Five supervised learning algorithms were implemented in R (version ≥ 4.3) using the *caret* framework: Multinomial Logistic Regression (*multinom*), Random Forest (*rf*); Support Vector Machine (radial kernel, *svmRadial*), k-Nearest Neighbors (*knn*) and Decision Tree (*rpart*). Models were first splited into trained dataset (80%) and test dataset (20%). And then the trained dataset was proceeded under five-fold cross-validation and models were fine-tuned by using grid-search on a range of hyperparameters. Each trained model was saved as an .rds file and the best model was used for downstream deployment in the Shiny App.

### 2.4 Evaluation and interpretability

Model performance was assessed using accuracy, F1 score, balanced accuracy and area under the ROC curve (AUC). For interpretability, SHAP analysis (via *kernelshap* and *shapviz* packages) quantified the contribution of each fluorescence parameter to model predictions. Global importance (bar plots) and individual variability (beeswarm plots) were generated for the best-performing model.

### 2.5 Implementation and reproducibility

The entire workflow was executed on a standard workstation-MacBook (≥ 4 cores, 8 GB RAM). Source code (*yield_predictor_classification*.*R and shinyapp*.*R*) and example datasets (*feature_data*.*txt* and *target_variables*.*txt*) are available upon request or through the GitHub repository (https://github.com/zx0223winner/YieldLearn).

## 3 Results

We first explored the distribution of five physiological measurements (Fo, Fm, Fv/Fm, PI Inst., and Area) by Hansatech Pocket PEA in July and August (Figure 1). Both months have the similar average observations but the data of July turns to have more outliers than in August. We then explored the correlations of physiological measurements and agronomic traits, those relatively low correlations parameters were chosen for the training analysis (Figure 2, Figure 3, Figure S1). Then we chose five supervised ML models which can perform well on relatively small datasets: Multinomial Logistic Regression (*multinom*), Random Forest (*rf*); Support Vector Machine (*svmRadial*), k-Nearest Neighbors (*knn*) and Decision Tree (*rpart*). Across all agronomic traits, classification performance ranged from 0.59 to 0.72 in accuracy, with AUC values ranged from 0.6 to 0.75 in AUC (Figure 3 and 4, Figure 5). Both the Random Forest model and Decision Tree work well with nonlinear relationship achieved the best predictive accuracy for Yield_class (accuracy = 0.562), while the Support Vector Machine showed competitive results for the others including Booting_class (accuracy = 0.594, AUC = 0.737), Height_class (accuracy = 0.594, AUC = 0.512) and Lodging_class (accuracy = 0.562, AUC = 0.575) and Maturity_class (accuracy = 0.576, AUC = 0.594) (See more details in Table 1). The hyperparameters for the best model can be found in Supplementary Table S1. It should be noted that the physiological measurements from August were used here. Since the human error and environmental bias can impact the measurement, so we did not merge the July and August observations together.

**Table 1.**
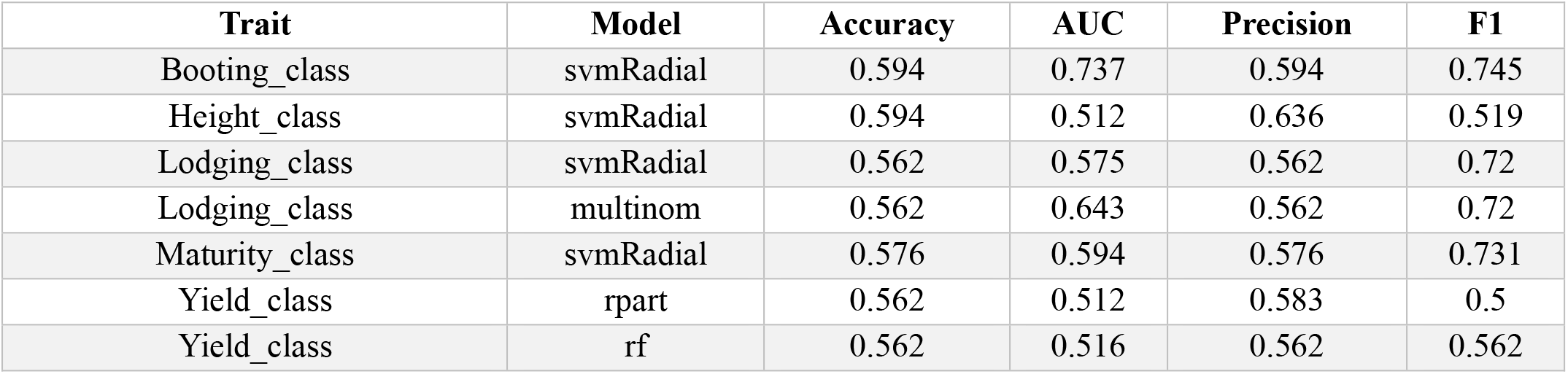
The ML models performance across different traits. Random Forest (rf); Support Vector Machine (radial kernel, svmRadial), and Decision Tree (rpart).

**Figure 1:**
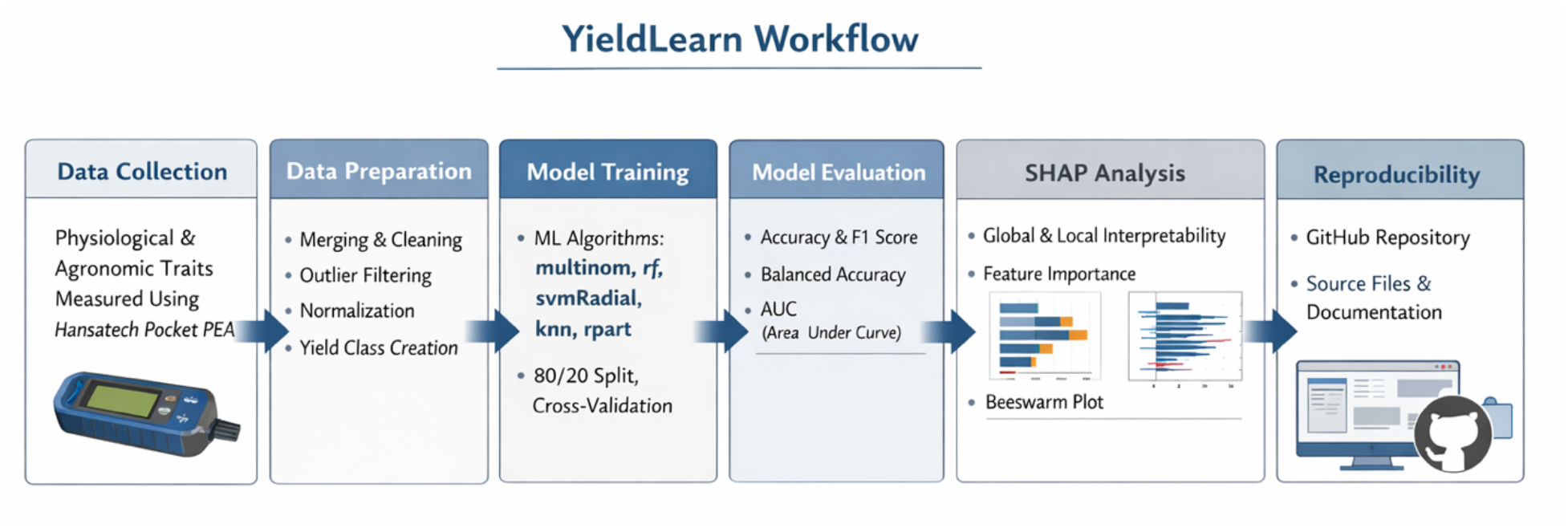
Workflow of the YieldLearn pipeline for predicting yield-related traits from physiological indicators. Multinomial Logistic Regression (multinom), Random Forest (rf); Support Vector Machine (radial kernel, svmRadial), k-Nearest Neighbors (knn) and Decision Tree (rpart).

**Figure 2:**
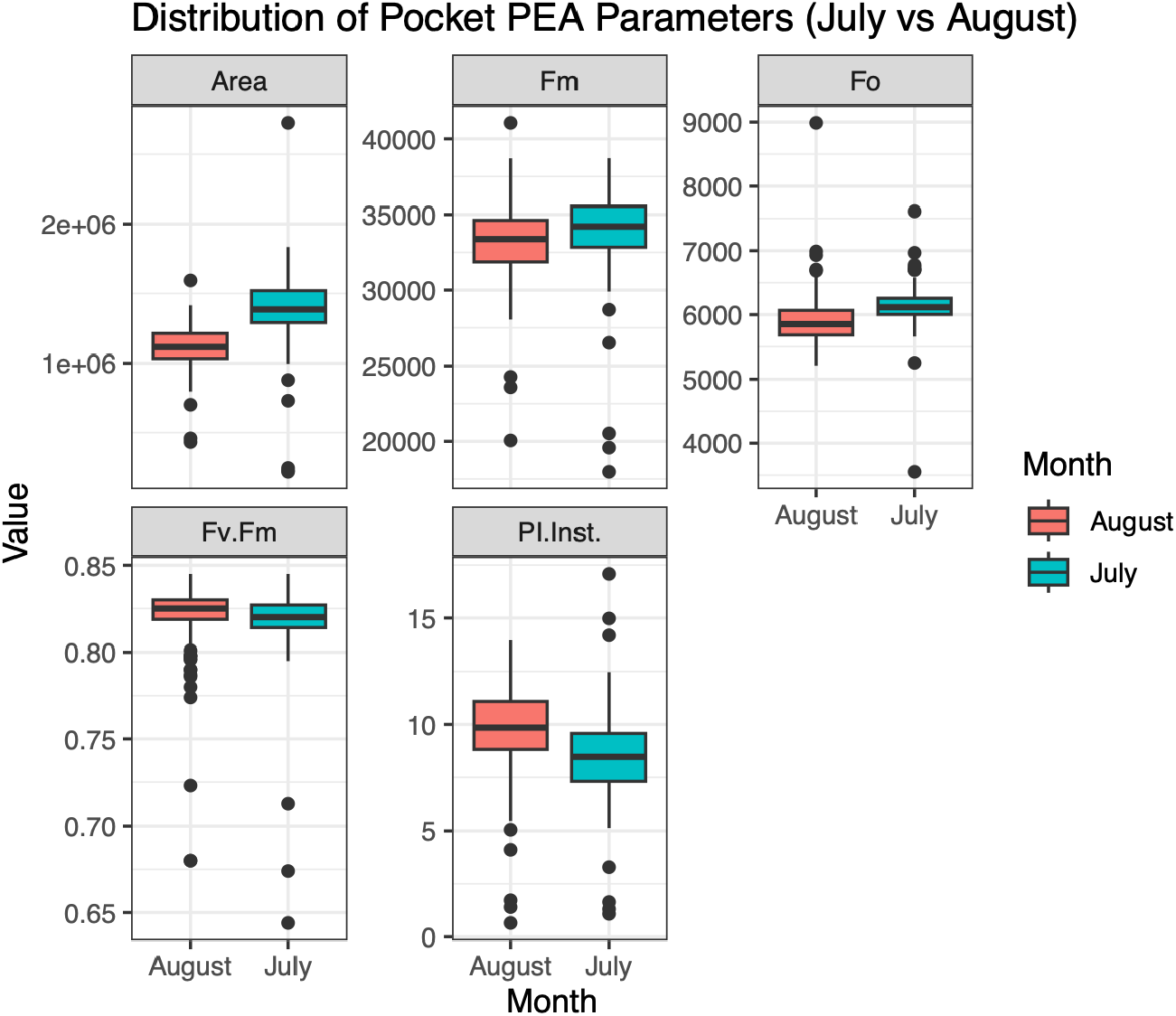
Distribution of physiological indicators (Fo, Fm, Fv/Fm, PI Inst., Area) measured in July and August using the Hansatech Pocket PEA.

**Figure 3:**
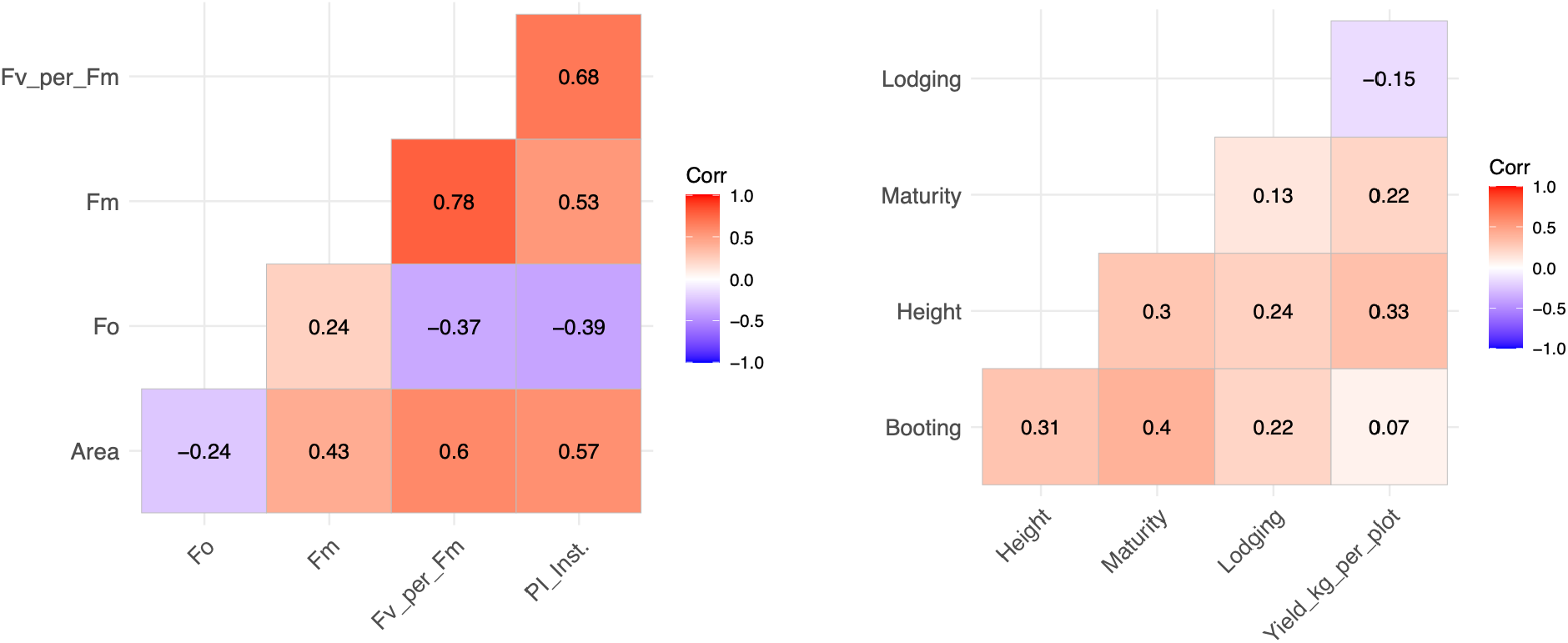
Correlation matrix comparing (left) fluorescence parameters (Fo, Fm, Fv/Fm, PI Inst., Area) and (right) agronomic traits (Booting, Height, Maturity, Lodging, Yield kg/plot) independently.

**Figure 4:**
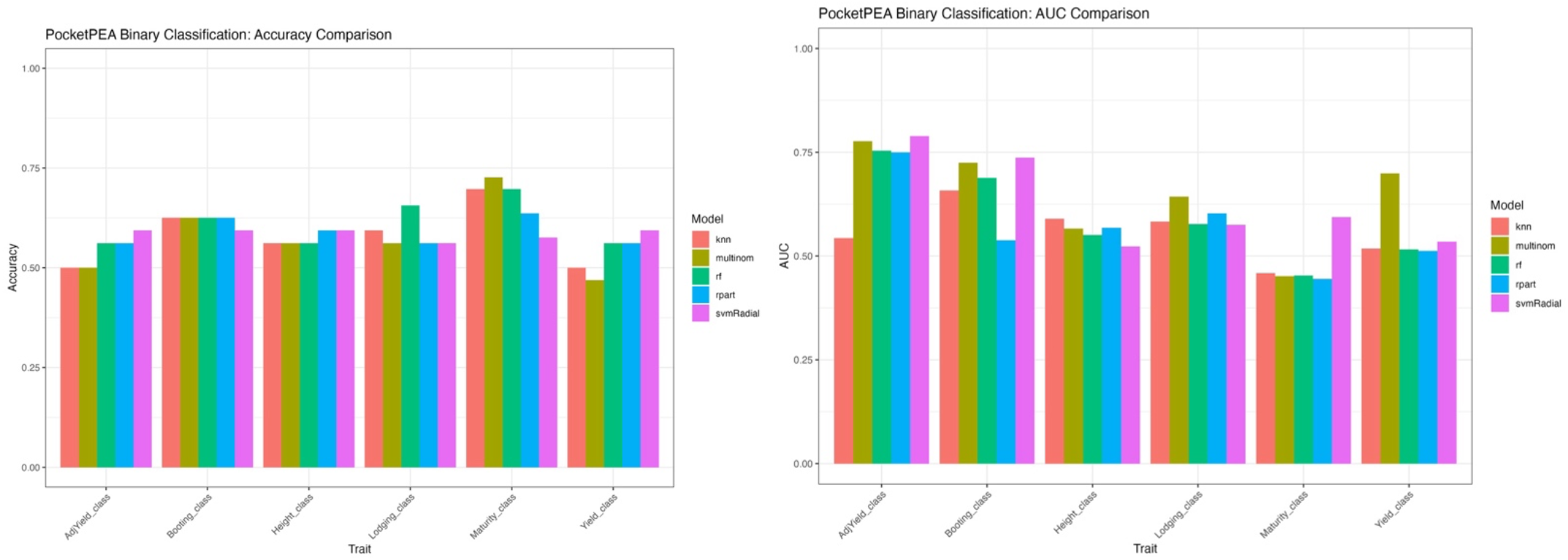
Visualization of classification (left) accuracy and (right)AUC for five supervised ML models (multinom, rf, svmRadial, knn, rpart) across agronomic traits: Booting, Height, Maturity, Lodging, and Yield classes.

**Figure 5:**
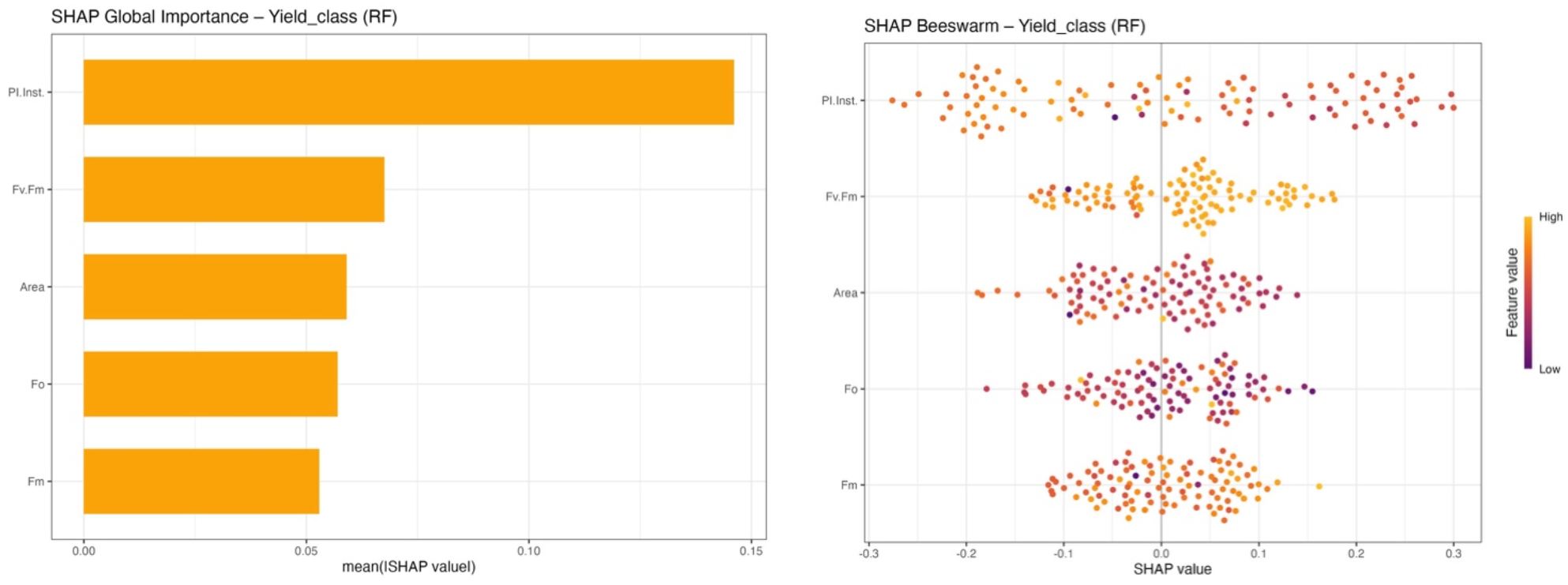
SHAP global importance plot (bar plot) and individual variability (beeswarm) for Yield_class. Bar plot showing the global SHAP feature importance values from the Random Forest model, indicating which fluorescence parameters contribute most to yield classification. Beeswarm plot illustrating sample-level SHAP value distribution for fluorescence predictors, highlighting how individual observations influence yield predictions in the Random Forest model.

Then SHAP visualizations were used to confirm the physiological relevance of fluorescence predictors. Global importance (bar plots) and individual variability (beeswarm plots) were generated for the random forest model on Yield_class (Figure 5). High Fv/Fm and PI Inst. values were positively associated with high yield, reflecting superior PSII efficiency and electron transport. Elevated Fo values indicated stress conditions corresponding to lower yield potential. We then use the saved random forest model to deploy into the YieldLearn Shiny App. It can provide an interactive interface where users can input fluorescence readings, select a target trait, and visualize predicted probabilities of high yield alongside SHAP-based feature-importance graphs (Figure 6). This facilitates immediate interpretation of physiological contributions within the field environment.

**Figure 6:**
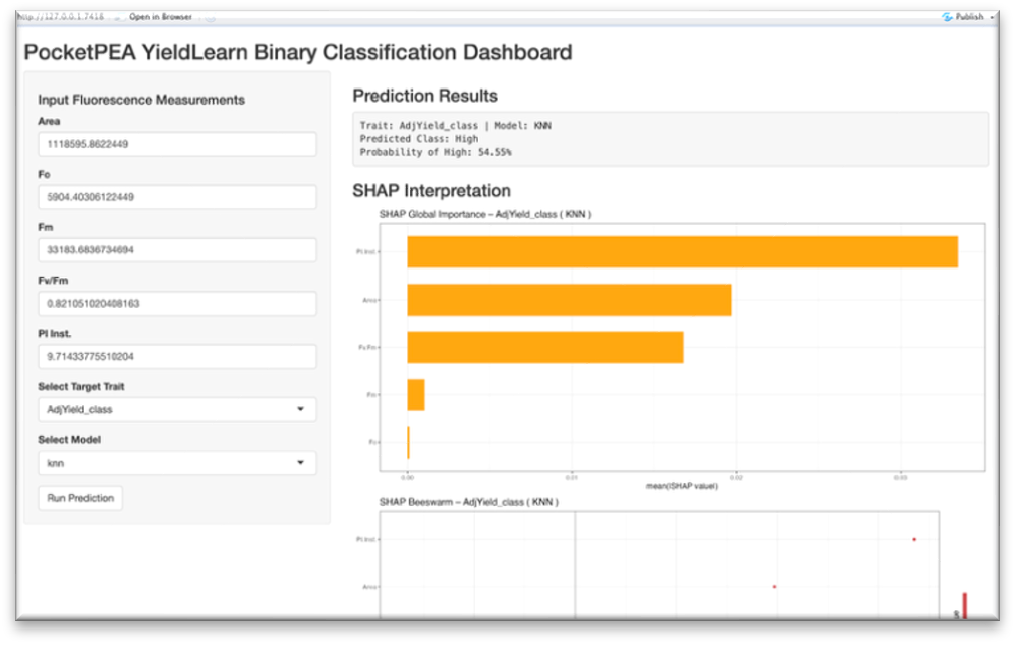
The interface of the YieldLearn Shiny app.

## 4 Discussion

This study demonstrates that fluorescence-based ML classification can distinguish high-and low-yield plots, outperforming traditional linear regression approaches. By capturing nonlinear interactions among physiological indicators, ensemble models such as Random Forest improve predictive capability even with modest sample sizes. Interpretability is a defining feature of YieldLearn. The integration of SHAP analysis provides transparent, quantitative explanations of how each fluorescence parameter influences model outcomes. The observed importance of Fv/Fm and PI Inst. aligns with known plant-physiological mechanisms: higher Fv/Fm indicates efficient energy conversion in PSII, while elevated PI Inst. reflects strong electron-transport capacity and stress resilience.

Despite encouraging results, limitations must be acknowledged. The relatively small dataset (165 plots) and lack of environmental covariates such as temperature, soil moisture, genotype or radiation restrict generalizability. Users should therefore apply YieldLearn with caution and validate its performance using larger, multi-environment datasets. Incorporating environmental and genotypic variables, as well as expanding to multi-class yield prediction, will enhance robustness and applicability. Nevertheless, this pipeline remains valuable as a reproducible baseline for physiological-trait modeling. YieldLearn and those of Gao et al. [3] demonstrate that photosynthesis-related parameters constitute a promising frontier for yield prediction—whether approached through genomic models or machine-learning pipelines. It provides a transparent starting point for integrating fluorescence data into digital phenotyping workflows, guiding experimental design, and informing future model development across crops and environments.

## Conclusion

YieldLearn establishes an interpretable, reproducible framework for estimating yield-related traits from physiological indicators. By combining multiple machine-learning algorithms with SHAP-based feature attribution and a user-friendly Shiny App, the pipeline bridges data analytics and biological insight. While current limitations in sample size and environmental scope constrain predictive generalization, the approach offers a strong foundation for future, large-scale, multi-factor modeling in digital agriculture.

**Figure S1:**
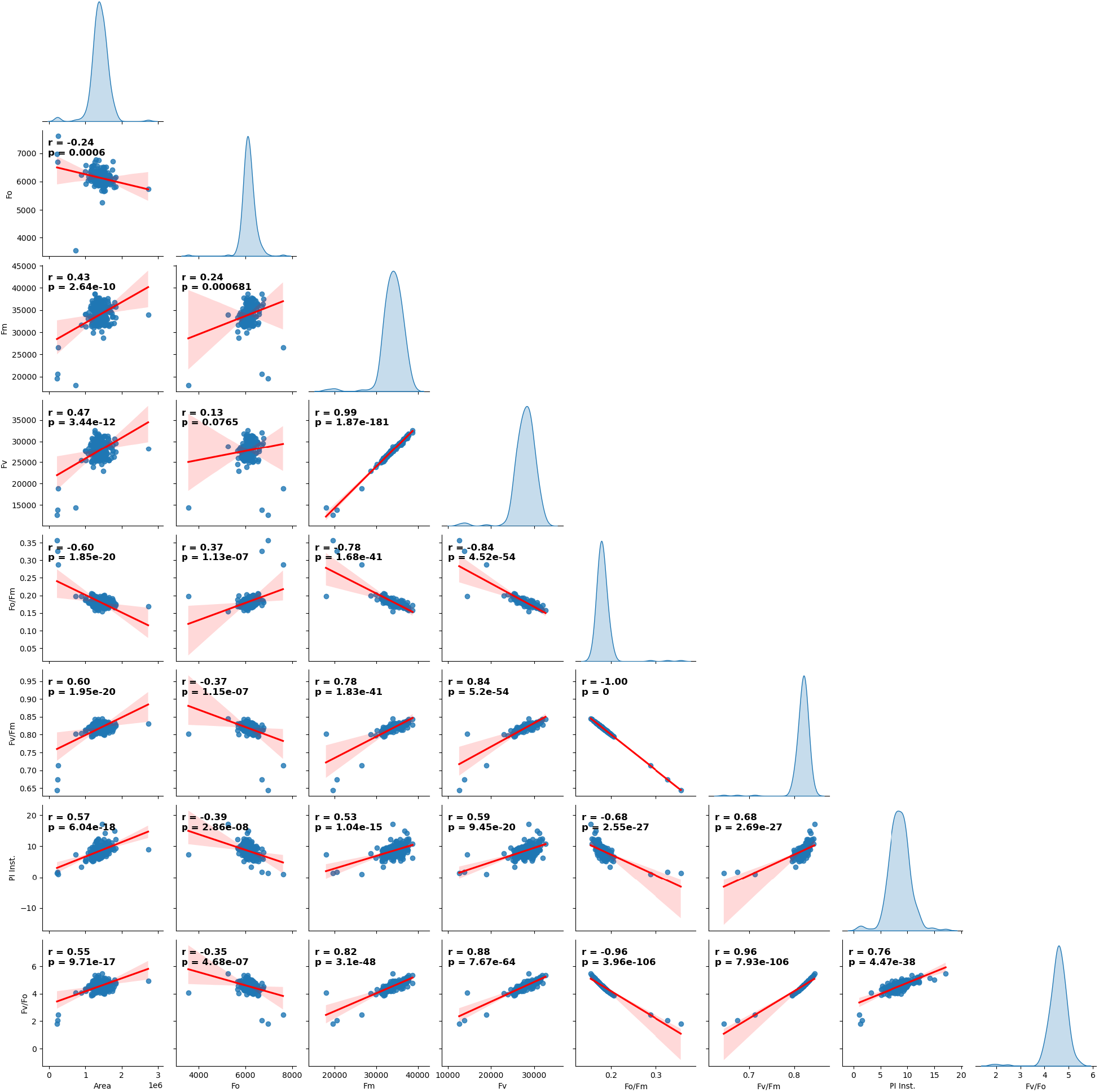
Supplementary visualizations supporting the correlation analyses of physiological indicators.

**Table S1:** Hyperparameters for the main model selection and fine-tuning.

| Trait | Model | mtry | sigma | C | cp |
| --- | --- | --- | --- | --- | --- |
| Booting_class | svmRadial | - | 0.001 | 0.500 | - |
| Height_class | svmRadial | - | 0.100 | 1 | - |
| Lodging_class | svmRadial | - | 0.010 | 0.500 | - |
| Maturity_class | svmRadial | - | 0.001 | 1 | - |
| Yield_class | rpart | - | - | - | 0.044 |
| Yield_class | rf | 3 | - | - | - |

## Acknowledgments

The authors gratefully acknowledge the support from the ACRD-NRC (Aquatic and Crop Resource Development Research Centre in National Research Council Canda)

## Author Contributions

The study was conceptualized by XZ. XZ wrote the initial draft. HS and PA collected the experimental data. All authors commented to produce the manuscript for peer review.

